## Supplementary Data for "YmoA functions as a molecular stress sensor in *Yersinia*"

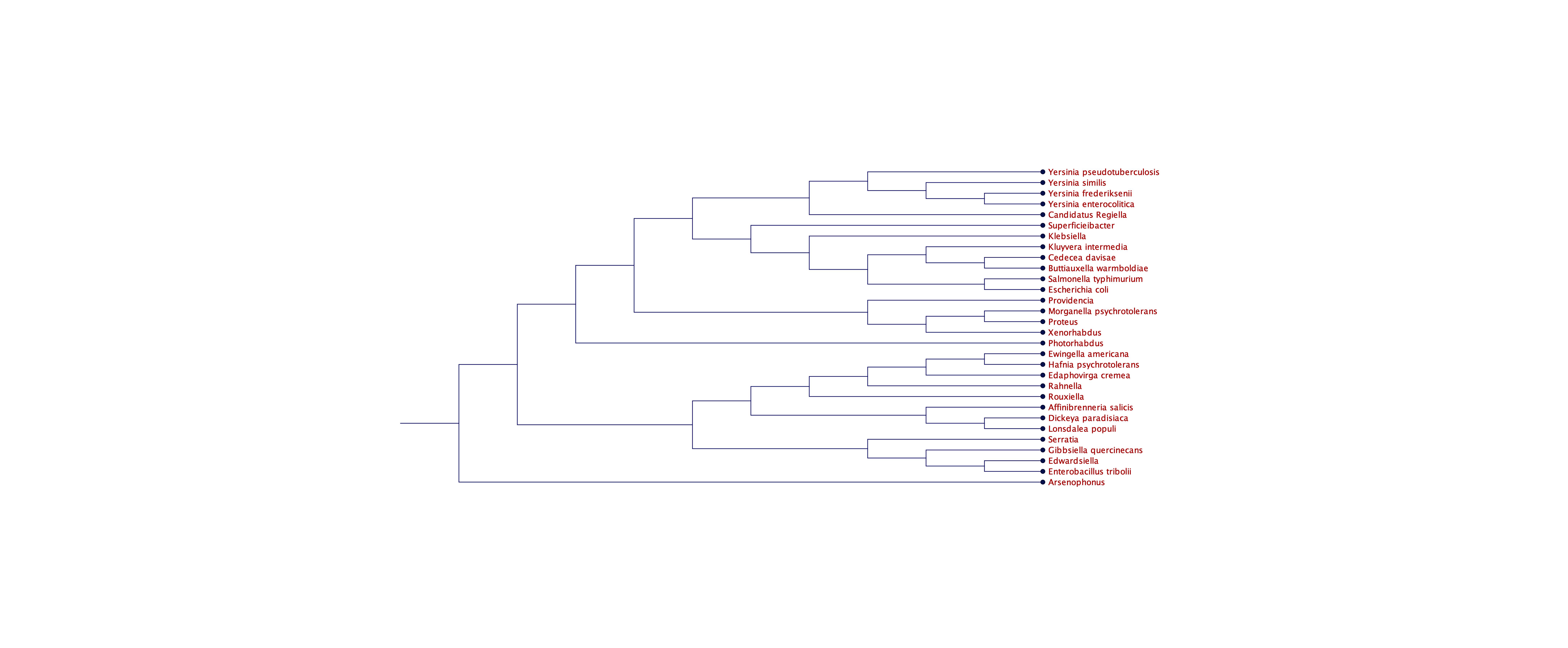

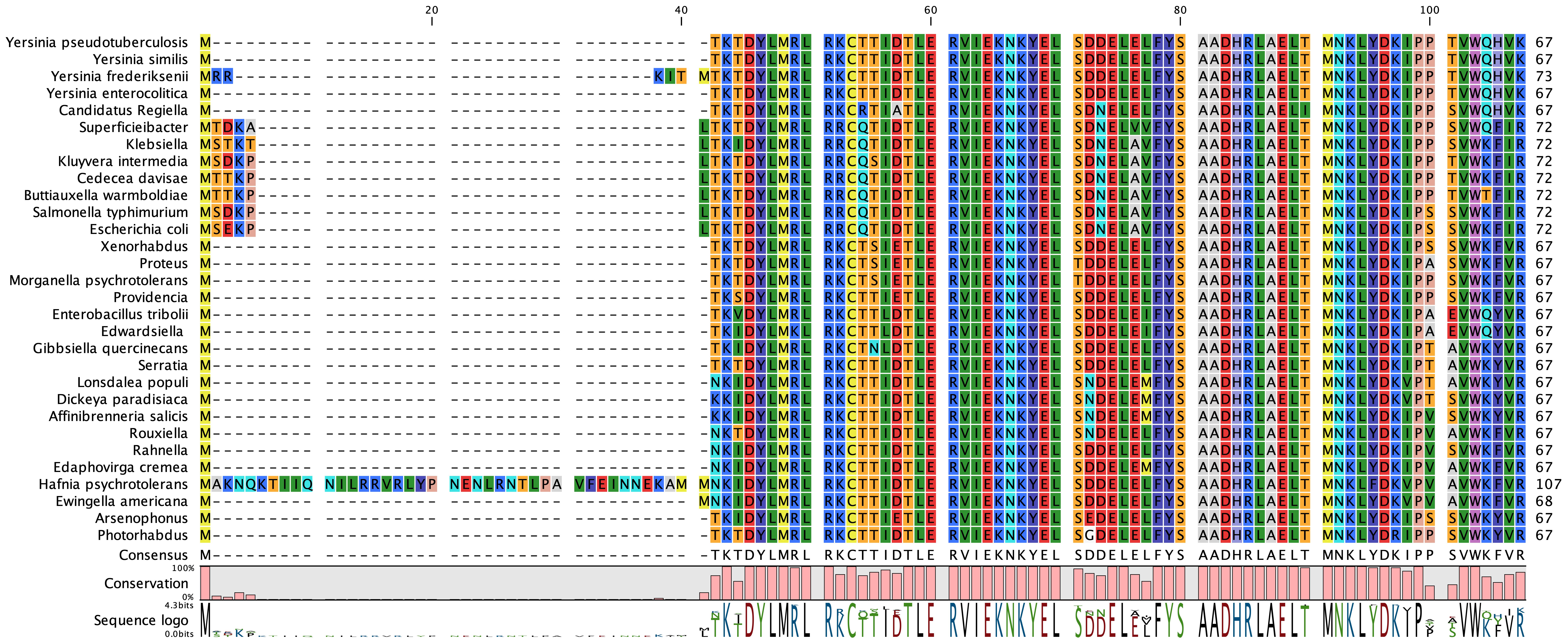

**Figure S1. YmoA/Hha is a well conserved family of protein.** Amino acids sequence alignment of YmoA/Hha gene from 30 species.

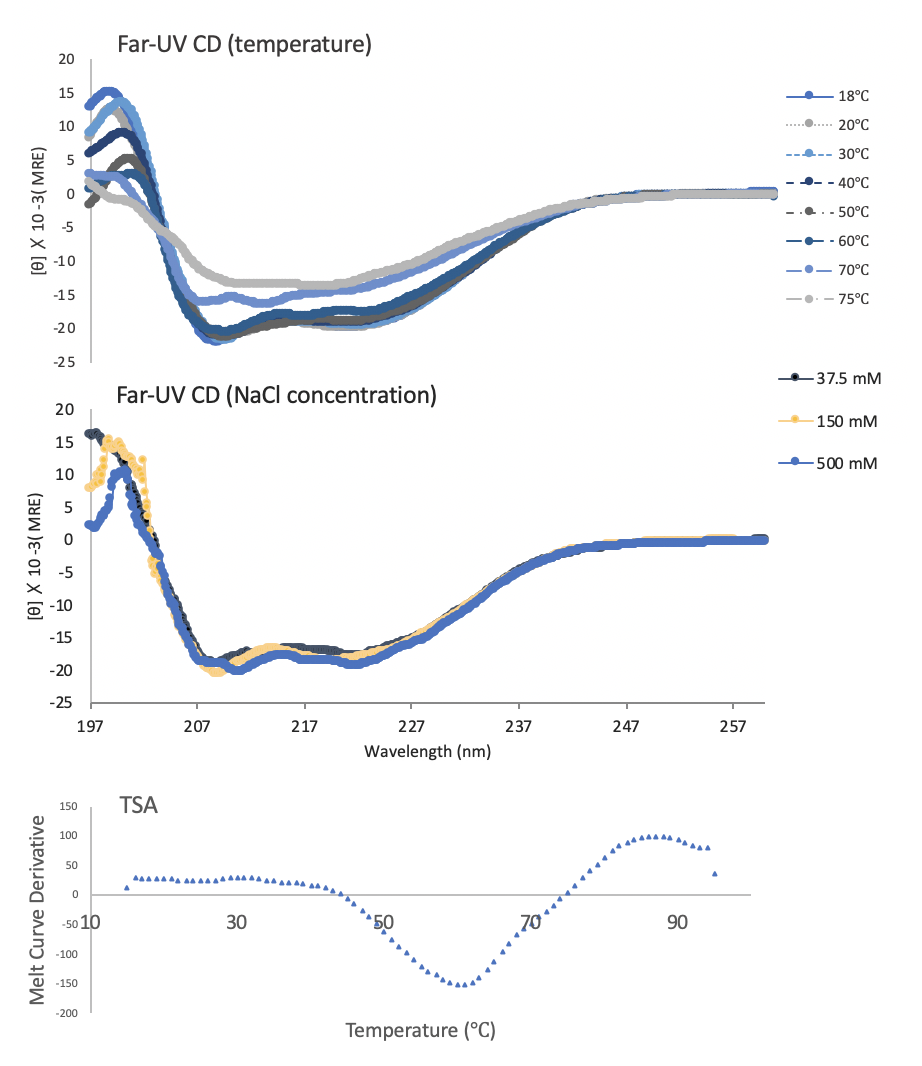

**a**

**b**

**Figure S2. YmoA is well folded and stable.** Both far-UV CD and thermal shift assay experiments confirmed that YmoA is folded, and stable at biological temperatures. **a. upper,** CD spectra show YmoA continuously loses the second structure over the increase of temperature from 18 to 75 °C. **a. lower,** YmoA possessed the same folding throughout the NaCl titration from 37.5 mM to 500 mM. **b.** Thermal shift assay also shows YmoA is folded below 50 °C and starts to unfold drastically at 60 °C.

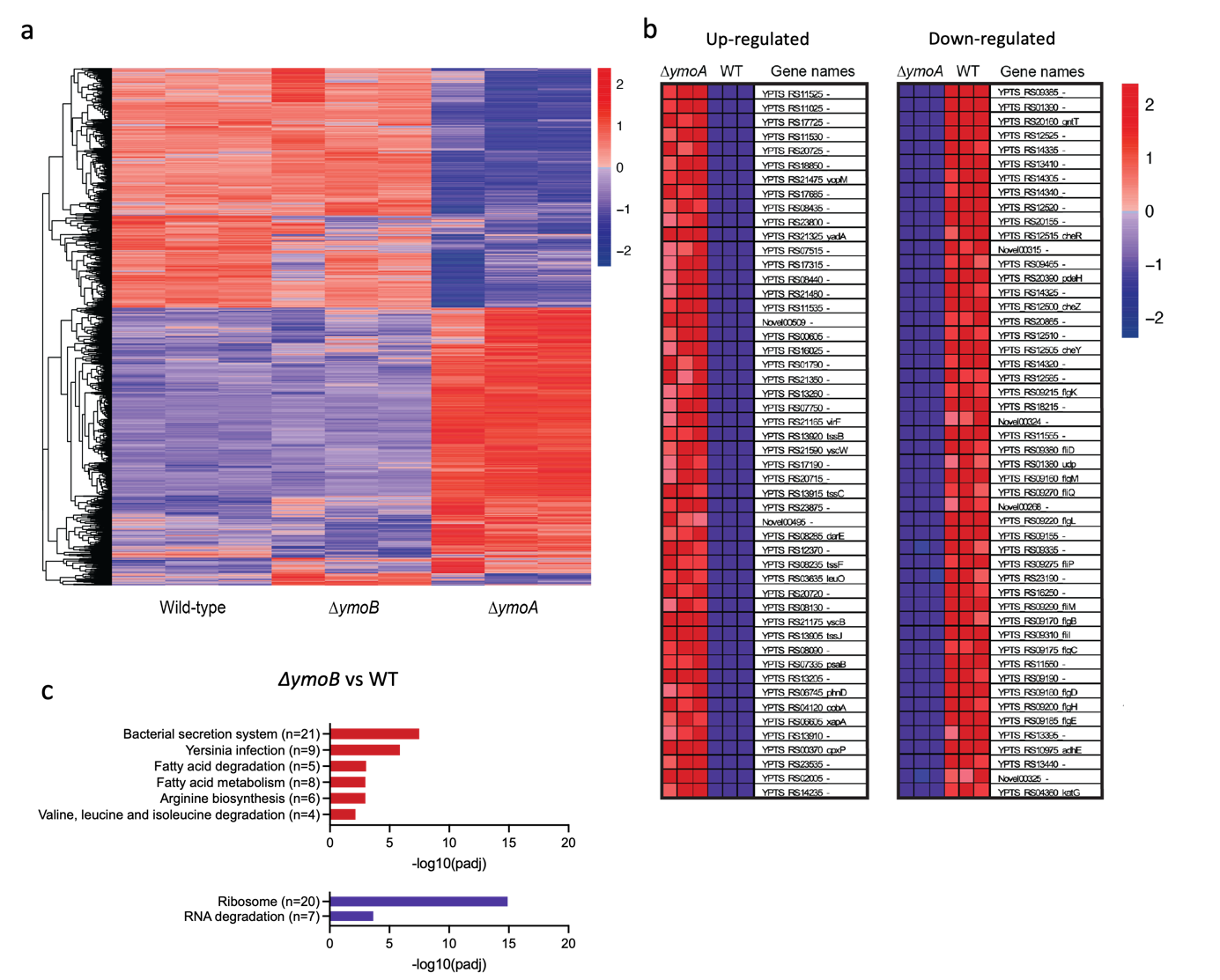

**Figure S3. RNA-seq analysis demonstrates up-regulation of T3SS genes and down-regulation of flagellar genes in a mutant ymoA. a-b.** Clustering heat map of genes**.** Red shows genes with high expression levels, purple shows genes with low expression levels. The red to blue color range represents the log_2_ (FPKM+1) value from large to small. **a.** Overall results of FPKM cluster analysis, clustered using the log_2_ (FPKM+1) value. **b.** Genes with high phenotype correlation, genes in descending order by the correlation of phenotypes according to signal2noise. Left panel, 20 most up-regulated genes. Right panel, 20 most down-regulated genes. **c.** Significantly enriched terms in the GO enrichment analysis. Upregulated genes are shown on the top in red and downregulated genes are shown on bottom in purple.

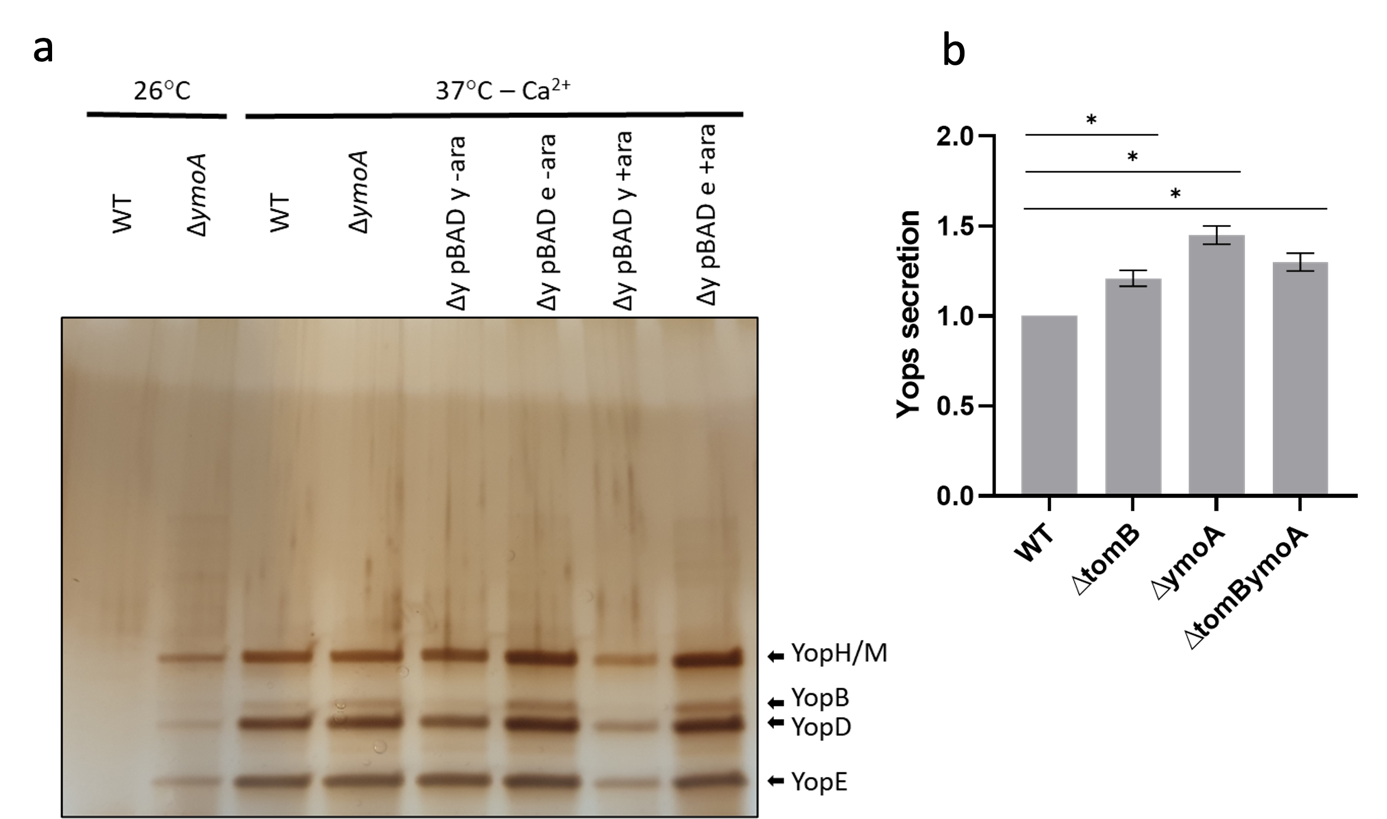

**Figure S4: Yop secretion.** a. Yops secretion of trans-complemented ∆*ymoA*. The same samples of bacterial supernatants as in Figure 2e were run on a SDS-page and silver-stained subsequently (n=1). b. Yop secretion of Figure 2e was quantified at T3SS inductive conditions (37°C -Ca2+) as the sum of YopE, YopD and YopH/M bands for each sample, normalized against the wild-type. The data shows the mean and SEM (n=3). Statistical analysis was done with a One-sample t-test (* p ≤ 0.05).

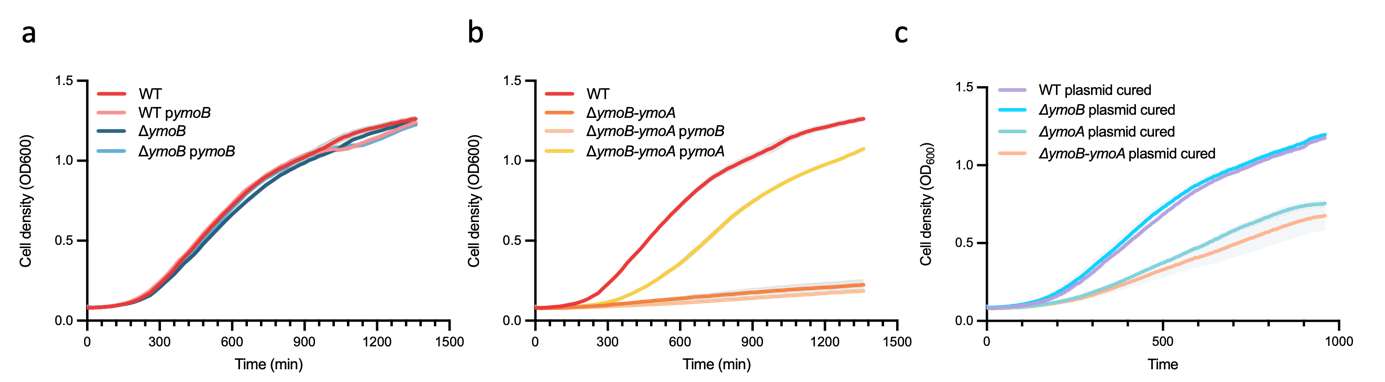

**Figure S5. Most of *ymoA* deletion fitness cost comes from the plasmid and *ymoB* has no effect on bacterial fitness. a-c.** Strains were grown at 26°C for 5h. Growth were determined by measuring the OD_600_. The data represent the mean ± SD (n=3).

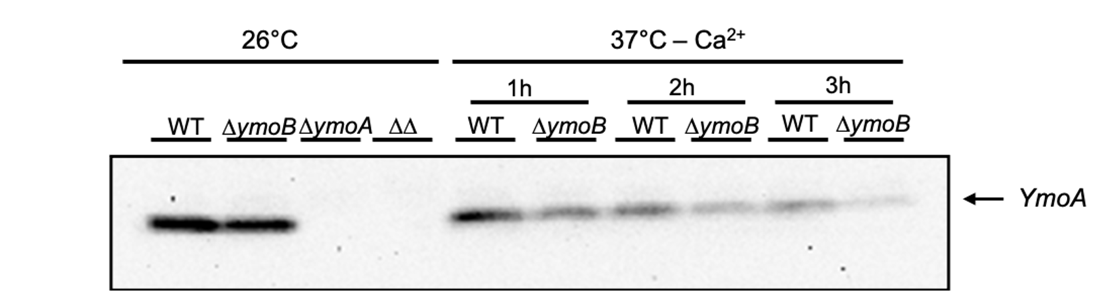

**Figure S6. YmoB reduces YmoA degradation at 37°C.** Western blots were performed on centrifuged pellet (whole cell) probed with anti-YmoA antibodies. Strains were grown at 26°C for 2h, subsequentially shifted to either 26°C or in T3SS inductive conditions: 37°C with Ca^2+^, and incubated for 1 h. Shown is one representative Western Blot from one out of 3 biological replicates.

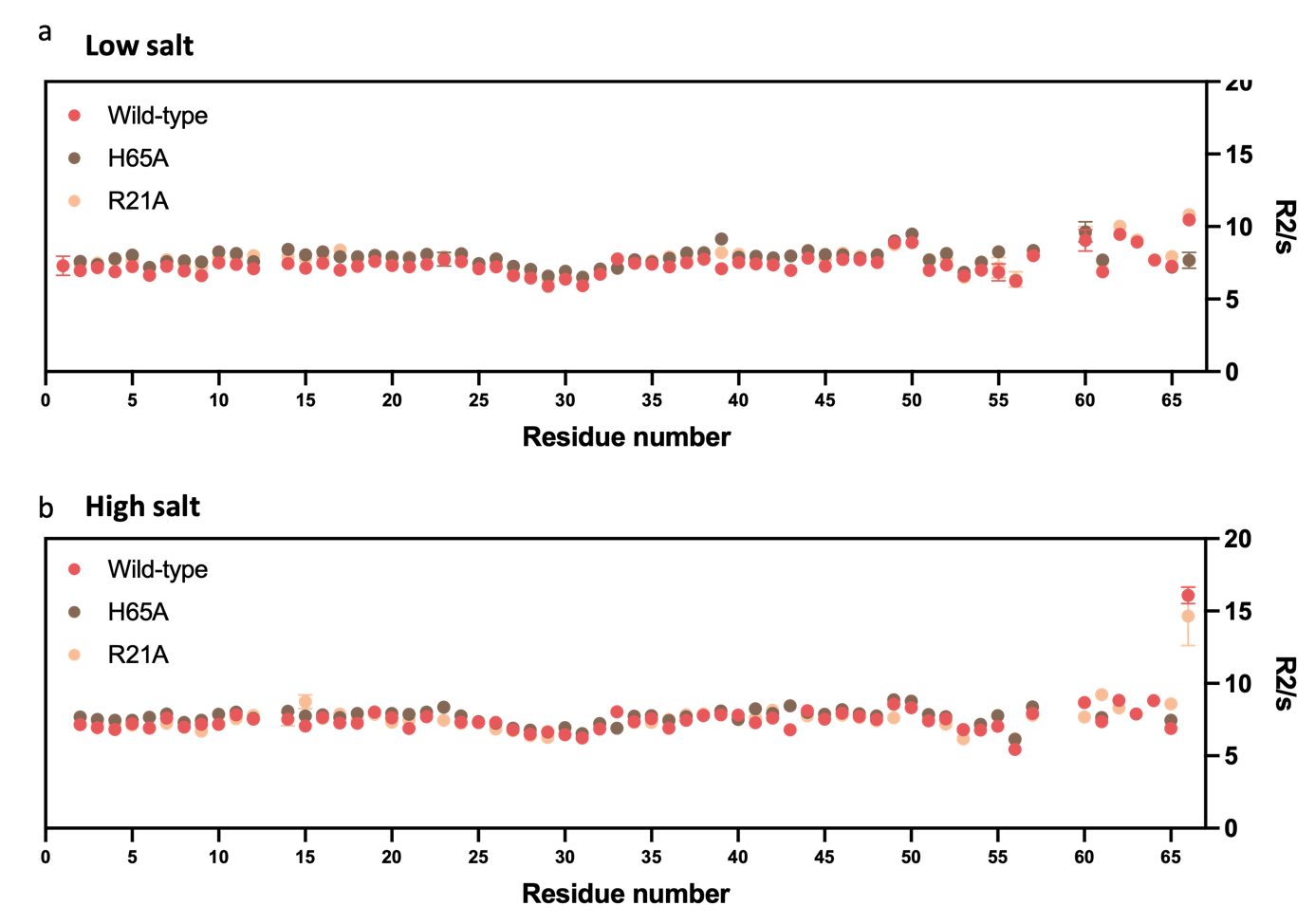

**Figure S7. High salt broadens YmoA Lys-67 relaxation time. a-b**. R2 relaxation measurement of the different YmoA residues in **a.** low salt 75mM or **b.** high salt 500mM concentration.

**Table S1**. Primers used for the generation of deletion mutants.

| **Name** | **Sequence** |
| --- | --- |
| YmoA fw | gcatgctagtaatgacaggccttctcctgcgg |
| YmoA rv | gccatacagtaggtggaattaaacgcatcaggtagtcagtttttgtcat |
| YmoB fw | gcatgcgtatttttaccccgatgactggatgattaatgaac |
| YmoB rv | gtacatacgtattccccgtcatgccgcttaggcgagtactcatccat |

**Table S2**. Primers used for protein purification.

| **Name** | **Sequence** |
| --- | --- |
| YmoA fw | atggatgagtactcgcctaagcggcat |
| YmoA_his_rv | ctaatgatgatgatgatgatgtttcacatgttg |
| YmoA_R21A fw | gatacgctagaagctgtaattgaaaaa |
| YmoA_R21A_rv | tttttcaattacagcttctagcgtatc |
| YmoA_H65A_fw | actgtatggcaagctgtgaaacat |
| YmoA_H65A_rv | atgtttcacagcttgccatacagt |
